# Fecal microbiota transplantation from individual with bipolar disorder and healthy control elicits distinct behaviors and metabolite profiles in mice

**DOI:** 10.1101/2023.11.16.566698

**Authors:** Grace Bukowski-Thall, Frederike T. Fellendorf, Sarah Gorkiewicz, Kenny Chi Kin Ip, Laura Schmidt, Marija Durdevic, Hansjörg Habisch, Sabrina Mörkl, Jolana Wagner-Skacel, Susanne A. Bengesser, Melanie Lenger, Nina Dalkner, Gregor Gorkiewicz, Christoph Högenauer, Tobias Madl, Christine Moissl-Eichinger, Aitak Farzi, Eva Z. Reininghaus

**Affiliations:** Division of Pharmacology, Otto Loewi Research Center, Medical University of Graz, Graz, Austria; Division of Psychiatry and Psychotherapeutic Medicine, Medical University Graz, Graz, Austria; Institute of Science and Technology Austria (ISTA), Klosterneuburg, Austria; Children’s Cancer Institute, UNSW Sydney, Sydney, New South Wales, Australia; Diagnostic & Research Institute of Hygiene, Microbiology and Environmental Medicine Medical University Graz, Graz, Austria; Austria; Core Facility Computational Bioanalytics, Center for Medical Research, Medical University of Graz, Graz, Austria; Institute of Pathology, Medical University of Graz, Graz, Austria; Otto Loewi Research Center, Division of Medicinal Chemistry, Graz, Austria; Division of Medical Psychology, Psychosomatics and Psychotherapy, Medical University of Graz, Graz, Austria; Department of Internal Medicine, Division of Gastroenterology and Hepatology, Medical University of Graz, Graz, Austria; BioTechMed-Graz, Graz, Austria

## Abstract

Bipolar disorder (BD) is a chronic mood disorder characterized by recurrent episodes of depression and (hypo-) mania. The gut microbiome is a potential avenue through which metabolic signaling, inflammatory pathways, environmental factors, and genetics influence BD pathogenesis via the gut-brain axis. Fecal microbiota transplantation (FMT) is a powerful translational tool for investigating the connections between the gut microbiome and BD, and there is evidence FMT can transfer affective symptoms of BD from humans to mice. In this study, we compared the behavior, gut-brain metabolomic profiles, and inflammatory marker expression in two groups of adult female C57BL/6J mice, one receiving FMT from a human donor with BD in a mixed episode ( HAM-D = 20, YMRS = 14) and another receiving FMT from a mentally healthy weight and age-matched control donor without BD (HAM-D and YMRS = 0). Here, we demonstrate that mice receiving FMT from individuals with BD had an increased abundance of Bacteroidota and decreased abundances of *Parabacteroides merdae* and *Akkermansia muciniphila* associated with altered levels of fecal metabolites, short-chain fatty acids, and related gut hormone expression relative to mice receiving control donor FMT. BD mice also exhibited differential regulation of several metabolites and inflammatory markers in the amygdala, with glycine being the most prominently affected. Furthermore, BD mice displayed increased anxiety-like behavior and decreased sociability, indicating that aspects of the behavioral phenotype of BD are transferable from humans to mice via FMT. Taken together, these findings implicate gut-brain signaling in the physiological and behavioral changes observed in our BD-FMT mouse model.

## Introduction

Bipolar disorder (BD) is a severe mood disorder characterized by recurrent episodes of depression, hypomania, and mania. Individuals with bipolar I (BDI) disorder undergo frequent depressive and manic episodes, and patients with bipolar II (BDII) disorder may experience a depressive or hypomanic episode without fully entering mania. BD is one of the most common psychiatric disorders worldwide [1,2] and is associated with increased mortality due to psychosocial consequences, suicidality, and psychiatric and somatic comorbidities [3,4]. Strengthening our understanding of the biological mechanisms behind BD is crucial for advancing treatment and improving patient outcomes. The psychiatric genomic consortium (PGC) revealed 64 genome-wide significant gene variants predisposing for BD [5], but genetics alone cannot explain the multifactorial phenotype of BD. Biopsychosocial interactions and gene-environment interactions play a crucial role in the outbreak and progression of BD [6,7]. There is also substantial evidence that the bidirectional signaling pathway known as the gut-brain axis is involved in BD pathogenesis [8]. Gut microbiota may influence BD by regulating neuroinflammatory systems [9], impacting gene expression [7,10], and producing metabolites and short-chain fatty acids (SCFAs) that influence CNS activity through vagus nerve interception [11–13]. BD patients also harbor different intestinal bacterial populations and have lower gut microbiome diversity than mentally healthy individuals [14]. Hence, disruptions in gut microbiome compositions by diet, medication, and other environmental factors can affect BD etiology [6,15].

Fecal microbiota transplantation (FMT) is a promising translational tool for BD research. One study found that depression-like behavior is transferable from humans to mice via FMT [16]. Mice receiving FMT from individuals with ADHD exhibited neurological changes and anxiety-related behaviors associated with ADHD [17]. Furthermore, FMT from a human donor with BD to mice affected the expression of TRANK1 [18], a gene whose down-regulation in the brain is associated with BD [19]. FMT also has therapeutic potential. FMT from a healthy human donor alleviated symptoms of alcohol-induced depression and anxiety in mice [20]. FMT from mentally healthy donors was an effective adjunctive treatment for depression for 4-8 weeks [21], and also improved symptoms of depression and mania for a male patient with BDII and ADHD [22].

Here, we used FMT to study the influence of gut-brain signaling on BD pathogenesis in a mouse model. Given the role of the gut-brain axis in BD, we hypothesized that mice receiving FMT from a human donor experiencing a mixed bipolar episode and mice with FMT from a mentally healthy age and weight-matched control donor would exhibit distinct emotional behaviors, inflammatory marker expression, and gut-brain metabolomic profiles.

## Materials and methods

### Animals

All experiments were approved by the ethical committee at the Federal Ministry of Education, Science and Research of the Republic of Austria (permit BMBWF-66.010/0047-V/3b/2019). Three cohorts of female C57BL/6J mice were bred in house and raised at a controlled temperature (22°C) and illumination (12:12 hr light-dark cycle, lights on at 07:00), and given access to water and a standard chow diet ad libitum.

### Donors and fecal sample processing

The study was conducted according to the guidelines of the Declaration of Helsinki and approved by the Institutional Review Board of the Medical University of Graz (protocol code 31-120 ex 18/19, date of approval: 27.03.2019). Informed consent was obtained from all subjects involved in the study. The Young Mania Rating Scale (YMRS) was used to assess the severity of manic symptoms [23], and the Hamilton Depression Scale for Depression (HAM-D) was used to assess the severity of depressive symptoms [24] in human donors. Fecal samples were collected from a female human donor in a mixed episode of bipolar disorder with a YMRS score of 14 and a HAM-D score of 20, as well as from a weight and age-matched healthy control with HAM-D and YMRS scores of 0.

The donor with BD was only on a very low dose of venlafaxine (37.5 mg/d) and refused mood stabilizers, resembling an almost drug-native state. The patient gave informed consent to our longitudinal study before in euthymic state. Exclusion criteria for both groups included antibiotic or antifungal treatment within the previous month, regular intake of prebiotics or probiotics, acute or chronic somatic diseases or infections (including rheumatoid arthritis, systemic lupus erythematosus, neurodegenerative and neuroinflammatory disorders), severe alcohol or drug abuse, dementia, history of digestive diseases such as inflammatory bowel disease and irritable bowel syndrome, history of gastrointestinal surgery (other than appendectomy), current pregnancy or breast-feeding, history of eating disorder (anorexia, bulimia) in the last two years, and laxative abuse. Self-reporting revealed no difference in somatization.

Fecal samples were collected in a plastic container containing an anaerobic generator (AnaeroGenTM 2.5L, Thermo Fisher Scientific). 60 to 100 g of the samples were processed within 6 hours of collection in an anaerobic chamber (Whitley A85 Workstation, Don Whitley Scientific). The stool samples were diluted in 150 mL of anaerobic, reduced sterile phosphate-buffered saline (PBS), homogenized and filtered through a sieve to remove solid components from the sample. 10% glycerol was added and the sample was aliquoted into 4 mL anaerobic tubes (Anaerobic Hungate Culture Tubes, Chemglass Life Sciences) [25]. Samples were stored at −70°C until stool transfer.

### Fecal microbiota transplantation (FMT)

Three cohorts of 20-30-week-old female C57BL/6J mice were treated with antibiotics added to their drinking water (0.5 mg/mL neomycin, 1 mg/mL meropenem, and 0.3 mg/mL vancomycin) for 1 week to deplete their existing gut microbiome and promote the uptake of donor microbiota as previously described [26]. A day prior to FMT, mice were fasted to prevent fecal blockage in the colon during transplantation and were given normal tap water instead of antibiotics. While anesthetized with isoflurane, one group of mice received stool from the donor with BD and another group was transferred stool from the healthy control donor. Approximately 0.2 mL of human stool sample was administered rectally into each mouse. A few drops of stool were also placed in the cage and on a few chow pellets to promote further microbiota uptake.

### Open Field Test

Cohort 1 mice were subject to the open field test 6 days post FMT as previously described [26]. The open field apparatus was a 35 lux illuminated 50 × 50 × 40 cm opaque, gray plastic box. Mice were placed in one corner of the open field and their locomotion (distance, velocity, speed) was measured for 5 minutes using a video camera and “EthoVision” (Noldus, the Netherlands) software. For analysis, the area was divided into a 36 × 36 cm center zone and the surrounding periphery zone. Entry into either zone was defined by the mouse center point.

### Elevated plus maze

Mice in cohort 1 were subject to the elevated plus maze (EPM) 10 days post-FMT as previously described [26]. The EPM apparatus comprised two open arms (30 × 5 cm), two closed arms (30 × 5 x 15.5 cm), and a center platform (5 cm × 5 cm). The maze was raised 60 cm above the floor and illuminated 35 lux from above. Time spent in the open arm, number of entries to open arm, and total distance traveled were recorded over the course of 5 minutes with “EthoVision” (Noldus, the Netherlands) software.

### Light/dark box test

Mice in cohort 1 were subject to the light/dark box test (LDT) 14 days post FMT [26]. The light-dark box was divided into two sections by a partition containing a door (4.5 × 6 cm) (TSE Systems). The light compartment (18.5 × 21 cm) was made of transparent walls and brightly illuminated (350-400 lux), whereas the dark compartment (18.5 × 21 cm) was made of black acrylic walls (20 lux). Mice were placed in the light compartment facing the dark compartment opening, and locomotion and exploration were tracked via two external infrared frames, which recorded light beam interruptions (counts) during a 10-minute period. Data was analyzed with LabMaster software (TSE Systems, Bad Homburg, Germany).

### Social interaction test

Mice in cohort 2 were subject to the social interaction test (SIT) 10 days post FMT. The test was performed in an open field apparatus containing two wire cages divided by a wooden panel extending halfway down the box. The SIT consisted of a 5-minute habituation phase and a 5-minute test phase during which the test mouse was exposed to a stranger mouse from a different home cage. The cage placement (right or left) of the stranger mouse was alternated between sessions. The time the test mouse spent in the vicinity (3 cm radius) of the stranger mouse’s cage vs the time it spent in the vicinity of the empty cage was measured with “EthoVision” (Noldus, the Netherlands) software, using the nose point of the test mouse as a reference. Each test mouse was given a one-hour interval between the habituation phase and the stranger mouse phase.

### Splash test

Cohort 2 mice were subject to the splash test 11 days post FMT. Mice were placed in individual cages and sprayed with a 10% sucrose and tap water solution to promote grooming behavior, which was quantified by latency, duration, and counts of both head and total body grooming. Mice exhibiting depression-like behavior groom less frequently [26]. Mice were recorded for 5 minutes after spraying, and grooming behavior was manually tracked with “EthoVision” (Noldus, the Netherlands) software.

### LabMaster cages

Circadian activity and sucrose preference were recorded continuously from 7 to 14 days post FMT for BD and control mice in cohort 3 using the LabMaster system (TSE Systems, Bad Homburg, Germany) as previously described [27]. The set-up consisted of six 42 x 26.5 x 15 cm test cages with two external infrared frames and a lid attached to two weight transducers. The hardware sampling rate was 100 Hz at the infrared frames and 1 Hz at the drinking and feeding sensors. The sampling interval of the LabMaster software was 1 minute. The two weight transducers were used to measure food and water intake. One drinking bottle was filled with water and another with a 1% sucrose solution. To record locomotion, the two external infrared frames were placed 4.3 cm apart horizontally, with the lower frame 2.0 cm above the bottom of the cage. The bottom infrared frame recorded horizontal locomotion (ambulatory movements) of the mice, while the top infrared frame recorded vertical movements (rearing, exploration). Mouse activity was measured by counts of interruptions of the light beams of the infrared frames.

### Tissue extraction

Tissue extraction was performed as described [28]. Mice were deeply anesthetized with pentobarbital (150 mg/kg) via intraperitoneal injection (IP). Brains were dissected and immediately frozen for 5 seconds in 2-methylbutane on dry ice. The amygdalas were microdissected under a stereomicroscope (Bregma −0.58 to −2.54) on a −20 °C cold plate (Weinkauf Medizintechnik, Forchheim, Germany) [29]. RNase AWAY (Carl Roth, Karlsruhe, Germany) was used to clean dissection instruments before and in between uses. Microdissected brain samples were placed in micro packaging tubes with Precellys beads (Peqlab, Erlangen, Germany) and stored at −70 °C until further analysis. Colon contents and distal and proximal regions were frozen on dry ice and stored at −70°C.

### Microbiome analysis

Bacterial DNA was extracted from fecal contents with the QIAamp DNA Mini Kit (Qiagen, Hilden, Germany) according to the manufacturer’s instructions. PCR amplification and sequencing was performed by Novogene Europe (Cambridge, UK). The variable V3–V4 region of the bacterial 16S rRNA gene was amplified with PCR using oligonucleotide primers 341F: 5-CCTAYGGGRBGCASCAG and 806R: 5-GGACTACNNGGGTATCTAAT, while additionally including a molecular barcode sequence. PCR reactions were carried out using the Phusion High-Fidelity PCR Master Mix (New England Biolabs). The PCR products were selected by 2% agarose gel electrophoresis. Same amount of PCR products from each sample was pooled, end-repaired, A-tailed and further ligated with Illumina adapters. Libraries were sequenced on a paired-end Illumina platform (NovaSeq PE250) to generate 250 bp paired-end raw reads. Paired-end reads were assigned to samples based on their unique barcodes and trimmed by cutting off the barcode and primer sequences. Paired-end reads were merged using FLASH (V1.2.7) [30]. Quality filtering was performed according to the Qiime quality control process [31] and chimeras were removed using UCHIME algorithm. Sequencing analyses were performed by Uparse software (Uparse v7.0.1090) [32] and sequences with ≥97% similarity were assigned to the same OTUs. Species annotation was performed by QIIME2 software using the annotation database SILVA138. Alpha diversity was calculated for observed species. The QIIME2 tool on the local Galaxy instance (https://galaxy.medunigraz.at/) was used to calculate beta diversity (weighted Unifrac) and predict bacterial biological pathways using PICRUst2.

### Metabolomics

Fecal and amygdala samples collected 13 days post FMT from cohort 1 were prepared for NMR spectroscopy measurements as previously described [33]. 50 to 100 mg of stool or brain tissue were mixed with 200 µl water and 400 µl methanol to inactivate and precipitate proteins. Remaining solids were lysed using a Precellys homogenizer (Bertin Technologies SAS, Montigny-le-Bretonneux, France), stored at −20 °C for 1 hour, followed by centrifugation for 30 min at 10,000 x g at 4 °C. Finally, the supernatants were lyophilized, resuspended in 500 μl NMR buffer (0.08 M Na_2_HPO_4_, 5 mM 3-(trimethylsilyl) propionic acid-2,2,3,3-d_4_ sodium salt (TSP), 0.04 (w/v) % NaN_3_ in D_2_O, with pH adjusted to 7.4 with HCl or NaOH, respectively), and transferred into 5-mm NMR tubes for measurement on the NMR instrument using the CPMG pulse sequence as described. Spectra pre-processing and data analysis have been carried out using Matlab® scripts (group of Prof. Jeremy Nicholson at Imperials College London). NMR data were imported to Matlab® vR2014b (Mathworks, Natick, Massachusetts, USA), regions around the water, TSP, and remaining methanol signals excluded, aligned, and corrected for sample metabolite dilution by probabilistic quotient normalization [34]. Reported values correspond to arbitrary units (A.U.) derived from the area under the peak being proportional to concentration.

### Quantitative polymerase chain reaction

RNA from the colon and amygdala was extracted with the RNeasy Tissue Mini Kit (Qiagen) and RNeasy Lipid Tissue Mini Kit (Qiagen), respectively. Aliquots of 1 µg RNA were reverse-transcribed with the High-Capacity cDNA Reverse Transcription kit (Applied Biosystems, Foster City, CA, USA). For relative quantification of mRNA, real time PCR was performed with the CFX384 Touch Real-Time PCR System (Biorad) using TaqMan inventoried gene expression assays for Tnf (Mm 00443258_m1), Il1b (Mm 00434228_m1), Il6 (Mm 00446190_m1), Ifng (Mm 01168134_m1), Ido1 (Mm 00492590_m1), Il10 (Mm 01288386_m1), Ccl2 (Mm 00441242_m1), Chil3 (Mm 00657889_mH), Iba1 (Mm 00479862_g1), Cd68 (Mm 03047343_m1), Arg1 (Mm 00475988_m1), Cldn1 (Mm 00516701_m1), Cldn5 (Mm 00727012_s1), Tjp1 (Mm 00493699_m1), Ocln (Mm 00500912_m1), Glra1 (Mm00445061_m1), Glra2 (Mm01168376_m1), Glra3 (Mm00475507_m1), Glra4 (Mm00501674_m1), Glrb (Mm00439140_m1), Gpr158 (Mm00558878_m1) and the TaqMan Gene Expression Master Mix (Life Technologies). All samples were measured as duplicates. Actb, Gapdh and Ppil3 were used as reference genes. Quantitative measurements of target gene levels relative to controls were performed with the 2-ΔΔCt method using the mean value of the control group as the calibrator [35].

### Statistics

Differences among mouse groups were assessed by Student’s t-test, Mann-Whitney, two-way ANOVA or repeated-measures ANOVA combined with Sidak post hoc analysis where appropriate. Pearson’s correlation was calculated for bacterial taxa that differed between BD and control mice versus behavioral parameters of the EPM followed by Bonferroni correction. Statistical analyses were performed with GraphPad Prism 9 (GraphPad Software, Inc CA, USA). *Metastats* was used for detecting differentially abundant microbial features [36]. Linear discriminant analysis (LDA) Effect Size (LEfSe) calculations implemented in Galaxy were performed to identify the predicted pathways that differentiate FMT effects. The statistical analysis of the metabolomics data was done using the web-based analysis platform “MetaboAnalyst” (https://www.metaboanalyst.ca/, Version 5.0, last visited 11/3/2023). Statistical significance was defined as p ≤ 05.

## Results

### Compositional and functional microbiome profiles of BD and control mice

BD and control FMT mice exhibited distinct compositional and functional microbiome profiles 10 days post FMT. Principal coordinate analysis (PCoA) of fecal 16S rRNA data demonstrated that the gut microbiomes of BD and control mice clustered differently, as did the microbiomes of their respective human donors (Fig. 1A). Full microbiome engraftment did not occur, which is unsurprising given innate differences between human and murine intestinal tracts. 16S rRNA sequencing of fecal samples showed notable distinctions in microbiome compositions between BD and control mice (Fig. 1B). Compared to controls, BD mice had a significantly higher abundance of the phylum Bacteroidota and significantly lower abundances of *Akkermansia muciniphila* and *Parabacteroides merdae* (Fig. 1E). Alpha-rarefaction curves approached saturation for both BD and control fecal samples, indicating sufficient sequencing depth to capture microbiome diversity, while no difference in alpha diversity was present between BD and control mice (Fig. 1C). Metagenomic predictions by PICRUSt2 identified differential regulation of several metabolic pathways such as glucose-1-phosphate degradation, S-adenosyl-L-methionine salvage I, and tetrapyrrole biosynthesis between BD and control mice (Fig. 1D).

**Fig 1.**
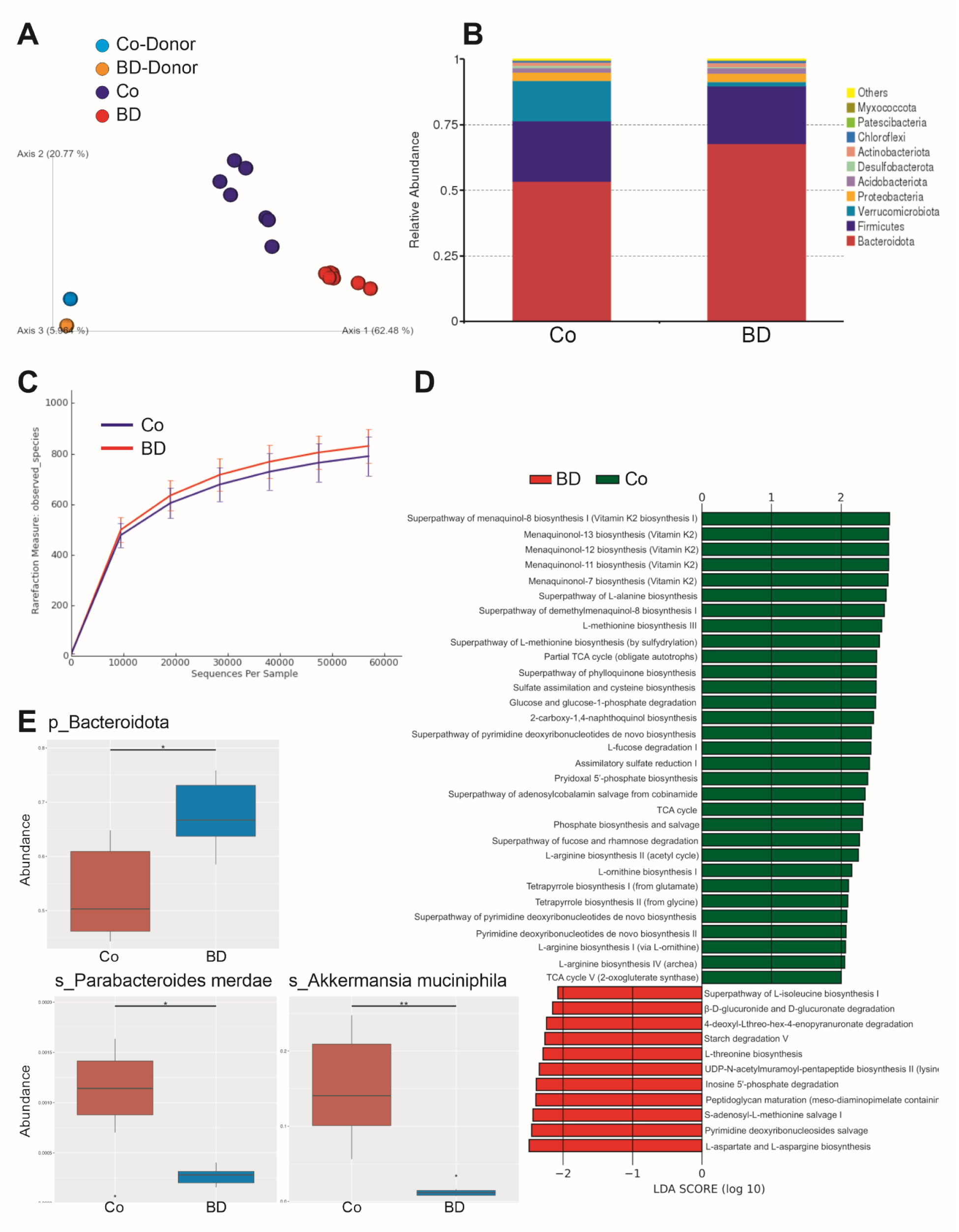
Compositional and functional microbiome profiles of female C57BL/6J mice 10 days after receiving FMT from a donor with BD or a control (Co) donor. PCoA plot of 16S rRNA gene data from feces of control donor (blue), donor with BD (orange), control mice (purple), and BD mice (red) (PERMANOVA, p < 0.001) (**A**). Bar plot showing 16S rRNA gene relative abundance of bacterial phyla in the feces of mice receiving FMT from an individual with BD and healthy control (**B**). Alpha-rarefaction curves (observed species) for both BD and control fecal samples show no differences in alpha diversity between BD and control mice (**C**). Enzyme Commission (EC) number pathways predicted by PICRUSt2 to be up-regulated in the microbiomes of BD (red) and control (green) mice selected by Linear discriminant analysis Effect Size (LEfSe) (**D**). Bar plot of fecal 16S rRNA sequencing results showing greater abundance of Bacteroidota phylum (q = 0.047) and lower abundance of *Parabacteroides merdae* (q = 0.034) and *Akkermansia muciniphila* species (q = 0.009) in BD mice relative to controls. Significant differences were analyzed using *Metastats* (**E**). BD n = 7, Co n = 7.

### Assessment of emotional behavior, circadian rhythm, and anhedonia in BD and control mice

BD-FMT mice exhibited more anxiety-like behavior and decreased sociability than mice with control FMT, but no differences in depression-like behavior. During the EPM 10 days post FMT, BD mice spent significantly less time in the open arms (Fig. 2E) and made significantly fewer entries to open arms (Fig. 2F). Time spent in the open arms also positively correlated with *Akkermansia muciniphila* in both BD and control groups (Fig. 2H). Traveling distance during the EPM was similar between BD and control mice (Fig. 2G). BD mice also displayed less sociability during the SIT 10 days post FMT. Control mice spent significantly more time in the vicinity of the stranger mouse cage than in the vicinity of the empty cage (Fig. 2J). By contrast, BD mice did not spend a significant amount of time exploring the stranger mouse over the empty cage (Fig. 2J). No significant differences between groups were measured during the OFT 6 days post FMT for time spent in the center (Fig. 2B), number of entries to the center (Fig. 2C), or total traveling distance (Fig. 2D). We measured no significant differences in time spent by each group in the light and dark compartments during the LDT 11 days post FMT (Fig. 2I). During the splash test 11 days post FMT, no significant difference in grooming duration was recorded between groups (Fig. 2K). No significant differences in circadian activity (Fig. 2L) or sucrose preference at any time point (Fig. 2M) were measured between BD and control mice 7 to 13 days post FMT.

**Fig 2.**
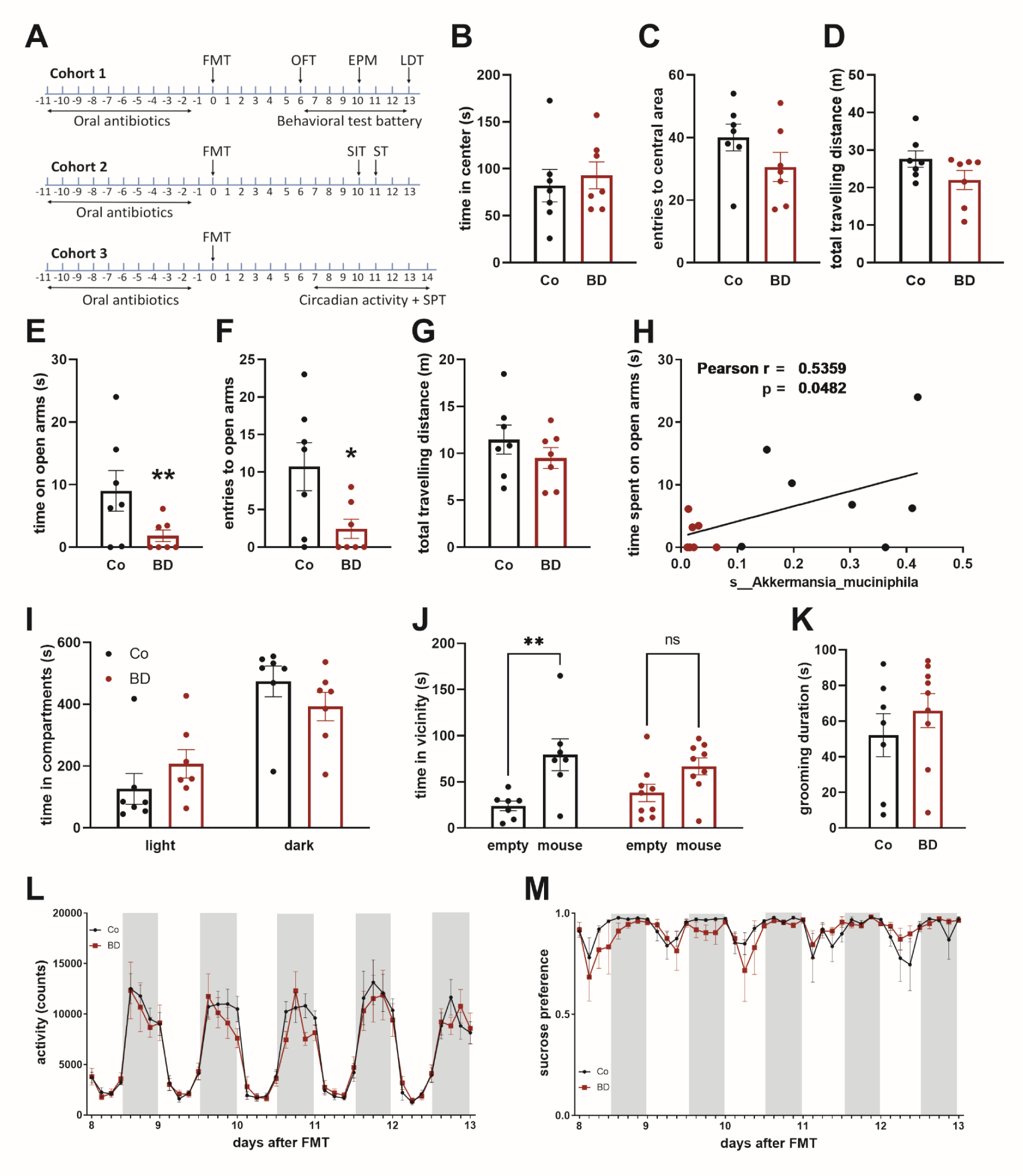
Assessment of emotional behavior, circadian rhythm, and anhedonia in 3 cohorts of mice receiving FMT from a donor with BD or a control (Co) donor. Experimental timeline, OFT = open field test, EPM = elevated plus maze, LDT = light/dark box test, SIT = social interaction test, ST = splash test, SPT = sucrose preference test (**A**). Comparison of time spent by BD and control mice in the center of open field (**B**), number of entries to the center of open field (**C**), and total traveling distance during OFT Comparison of time spent by BD and control mice in the open arms of EPM (p = 0.0079) (**E**), number of entries to open arms (p = 0.043) (**F**), and total traveling distance during EPM (**G**). Time spent in the open arms of EPM positively correlates with *Akkermansia muciniphila* relative abundance for both BD and control mice (Pearson r = 0.5359, p = 0.0482) (**H**). Relative time spent by BD and control mice in light and dark compartments of light/dark box (**I**). Total time spent in the vicinity of an empty cage vs time spent in the vicinity of a cage with a stranger mouse for BD (p > 0.05) and control mice (p = 0.0041) (**J**). Difference in grooming duration between BD and control mice during splash test (**K**). Number of activity counts for BD and control mice between 7 and 14 days post FMT (**L**). Percent sucrose preference for BD and control mice between 7 and 14 days post FMT (**M**). BD n = 7, Co n = 7. Data represent mean ± SEM. Significant differences between BD and control groups were analyzed using an unpaired t-test or 2-way ANOVA, *p ≤ 0.05, **p ≤ 0.01.

### Gut and brain metabolomic profiles of BD and control mice

BD and control mice displayed significantly different gut and brain metabolomic profiles 13 days post FMT. Metabolomic analysis revealed significant up-regulation of fecal valeric acid, fructose, glucose, and sucrose in BD mice relative to controls. Fecal aspartic acid, capric acid, xanthine, and hypoxanthine were significantly down-regulated in BD mice (Fig 3A). There were significantly lower levels of acetic acid and a notable, but not statistically significant reduction of butyric acid in the feces of BD mice. Fecal propionic and succinic acid levels were similar between BD and control groups (Fig. 3B). BD mice had significantly lower levels of fecal L-glutamic acid than control mice (Fig. 3C). Expression levels of gut hormones Peptide YY (*Pyy*) and preproglucagon (*Gcg)* were decreased in BD mice colons relative to controls (Fig. 3D). Given the increased anxiety-like behavior observed in BD mice, we focused on the amygdala for metabolomic analysis of the brain. Our results revealed significant down-regulation of glycine, glycerol, uridine, tryptophan, pyroglutamic acid, inosine, acetic acid, phosphorylcholine, choline, and methionine in the amygdalas of BD mice, with glycine being the most prominently affected. Conversely, adenosine monophosphate and inosinic acid were significantly up-regulated in the BD group (Fig. 3E).

**Fig 3.**
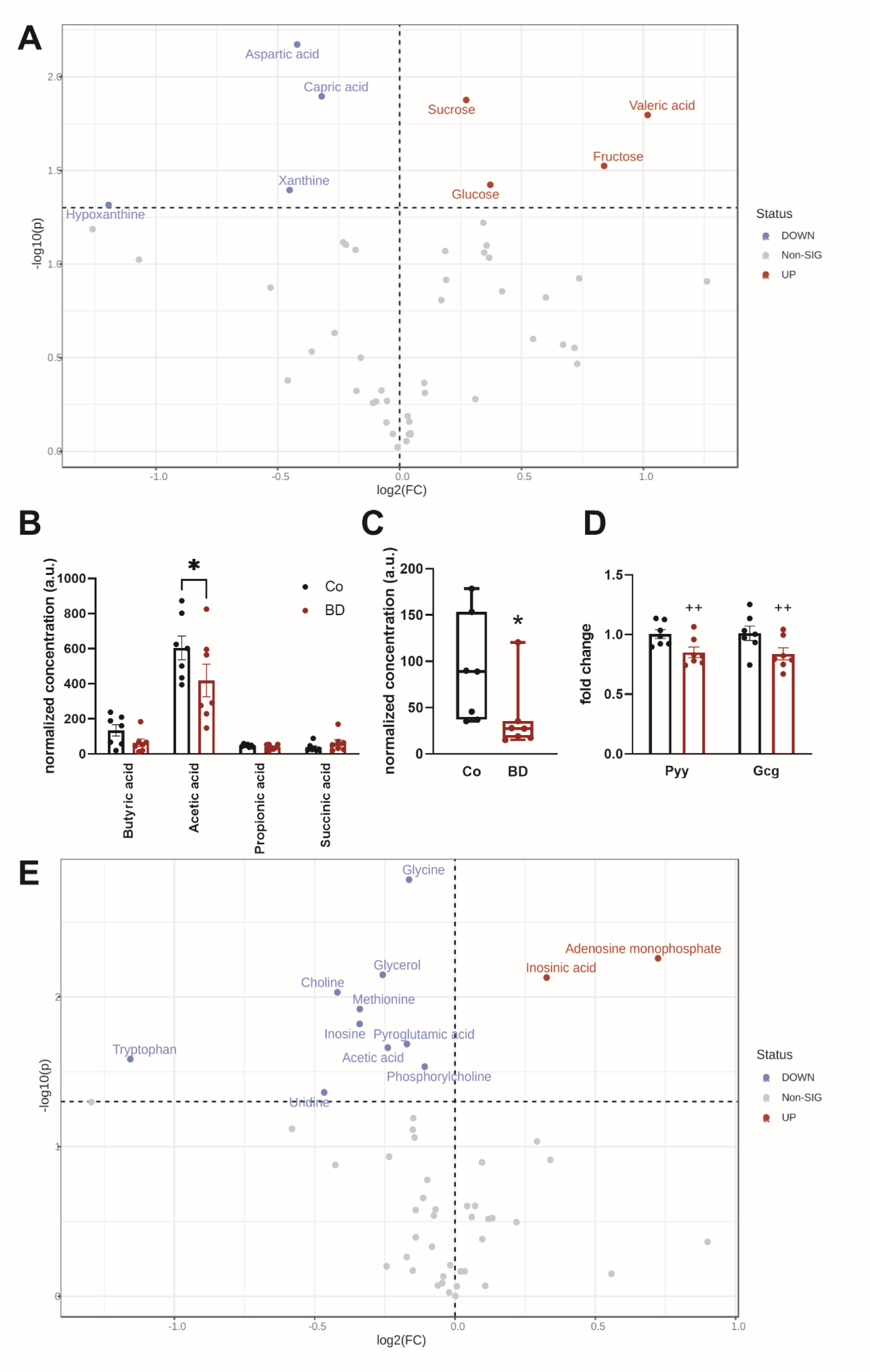
Metabolite and SCFA profiles of the feces and amygdalas of BD and control mice. Volcano plot representing the metabolomic profiles of the feces of BD mice plotted relative to control mice 13 days post FMT (**A**). Bar plots with relative levels of short-chain fatty acids acetic acid (2-way ANOVA, p = 0.017), butyric acid, propionic acid, and succinic acid in the feces of BD and control groups (**B**). Box plots comparing levels of fecal L-glutamic in BD mice relative to controls (Mann-Whitney, p = 0.011). (**C**). Bar plot showing expression of colonic mRNA expression of gut hormones *Pyy* and *Gcg* (2-way ANOVA main effect, p = 0.003) in BD and control mice (**D**). Volcano plot representing the metabolomic profiles of BD mice feces plotted relative to control mice For both volcano plots, the x-axis represents the log2 fold change (log2(FC)), indicating the magnitude of expression change. The y-axis represents the negative logarithm of the p-value (-log10(p)), indicating the statistical significance of the change. Metabolites with statistically significant upregulation are shown in red, while those with significant downregulation are in blue. Non-significant changes are in gray. The dashed horizontal line depicts the threshold for statistical significance (p < 0.05). BD n = 7, Co n = 7. For bar plots, data represent mean ± SEM, *p ≤ 0.05, **p ≤ 0.01, ^++^p ≤ 0.01.

### Cytokine, inflammatory marker, tight junction protein, and glycine receptor subunit expression in BD and control mice

Thirteen days post FMT, BD and control mice displayed similar levels of colonic (Fig. 4A) and amygdalar (Fig. 4C) *Ido-1* mRNA expression. Pro-inflammatory cytokine (*Tnf*, *Il1b*, *Il6*, *Ccl2*, *Ifng*) and anti-inflammatory cytokine (*Il10*) expression did not differ significantly between groups in both the colon (Fig. 4A) and amygdala (Fig. 4C). However, BD mice did exhibit a significant increase in amygdalar *Chil3* expression relative to controls (Fig. 4C). There were no significant differences between BD and control mice in the expression of tight junction protein genes *Cldn1*, *Cldn5*, *Tjp1*, and *Ocln* in both the colon (Fig. 4B) and the amygdala (Fig. 4D), nor was there differential expression of microglial markers *Iba1*, *Cd68*, *Cd86*, and *Arg1* (Fig. 4E). *Glra4* expression was significantly increased in the brains of BD mice relative to controls, but there were no significant differences in the expression of other glycine receptor subunits *Glra1*, *Glra2*, *Glra3*, *Glrb*, and *Gpr158* (Fig. 4F).

**Fig 4.**
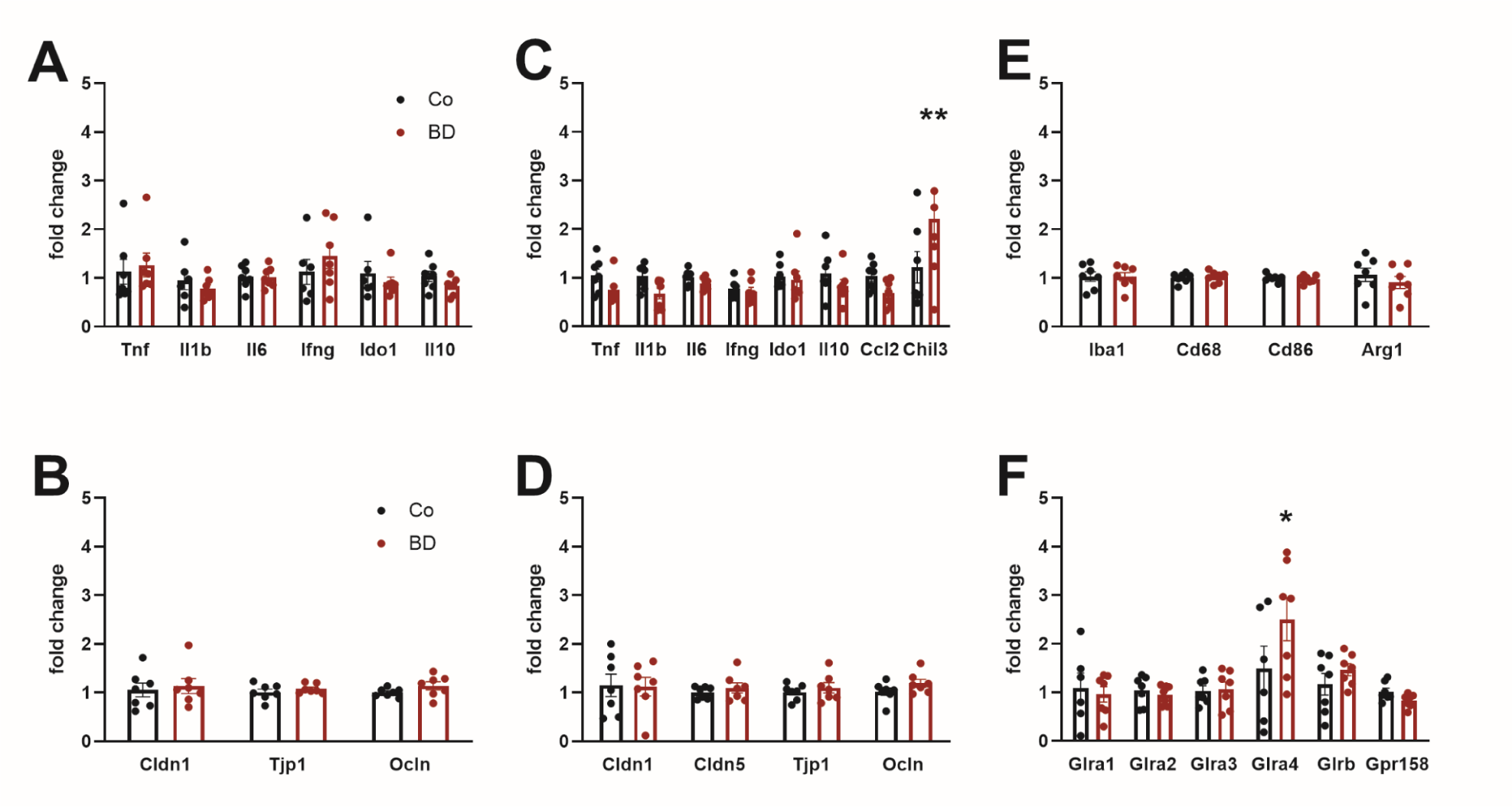
Quantitative polymerase chain reaction (qPCR) determined mRNA expression levels of tight junction proteins, cytokines, and other inflammatory markers in BD and control (Co) FMT mice groups. Bar plot showing that relative colonic mRNA expression of cytokines, inflammatory markers (**A**), and tight junction proteins (**B**) were not significantly different between BD and control mice. Bar plot showing up-regulated mRNA expression of *Chil3* in the amygdalas of BD mice relative to controls (p = 0.0047), but expression of other cytokines and inflammatory markers were not differently regulated (**C**). Bar plot showing that tight junction protein expression was similar between BD and control mice (**D**). Bar plot showing that amygdalar mRNA expression levels of glial markers in BD and control mice were not significantly different between BD and control groups (**E**). Bar plot showing that *Glra4* expression was significantly elevated in BD mice (p = 0.015), but other glycine receptor subunit proteins were not differentially expressed between groups (**F**). BD n = 7, Co n = 7. Data represent mean ± SEM. Significant differences between BD and control groups were analyzed using 2-way ANOVA, *p ≤ 0.05, **p ≤ 0.01.

## Discussion

Mice receiving FMT from a donor with BD exhibited altered microbiome compositions associated with differences in behavior and fecal and brain metabolomic profiles relative to mice with FMT from a mentally healthy control. These findings implicate the involvement of gut-brain signaling in the physiological and behavioral changes in our BD-FMT mouse model and provide a promising translational framework for understanding the mechanisms of BD.

Increased anxiety-like behavior and decreased sociability of BD mice suggest that these behavioral features are especially affected by gut-derived signals. The positive correlation observed between *A. muciniphila* and open arm visits in the EPM is supported by previous findings that *A. muciniphila* relative abundance negatively correlates with anxiety-like behavior in mice [37]. Furthermore, probiotic treatment with *A. muciniphila* has been shown to reduce anxiety-like behavior, motor degeneration, and learning memory deficits in mice [38–40]. A murine model of mania demonstrated hyperactivity, disrupted circadian rhythm, and decreased sucrose preference [41]. However, a lack of significant differences between BD and control FMT mice in these areas suggests that the manic and depressive symptoms of the BD donor were less transferable to mice via FMT.

Behavioral differences between BD and control mice are possibly linked to differential regulation of several gut and brain metabolites, short-chain fatty acids (SCFAs), and related gut hormone signaling. Reduced *A. muciniphila* in BD mice potentially lowered fecal acetic acid concentrations, since *A. muciniphila* produces acetic acid through intestinal mucin degradation [42]. Acetic acid was also decreased in the amygdala, suggesting a link to fecal acetic acid depletion. Elevated fecal valeric acid in BD mice possibly resulted from reduced *P. merdae,* which can catabolize branched-chain amino acids such as valine [43]. *P. merdae* also inversely correlates with depression in humans [44]. Interestingly, increased valeric acid and decreased acetic acid concentrations have been measured in fecal samples from children with autism [45], suggesting a connection between the regulation of these metabolites and neurodevelopmental disorders. Acetate supplementation has also rescued social deficits in an autistic mouse model, implicating decreased central acetic acid levels with impaired social interaction in BD mice [46].

Diminished acetic and butyric acid production in BD mice possibly reduced colonic expression of gut hormones *Gcg* and *Pyy* [47,48]. In mice, *Gcg* encodes for glucagon but also produces glucagon-like peptide-1 (GLP-1) [49], which is released from intestinal cells and can cross the blood-brain barrier [50]. GLP-1 attenuated symptoms of mania in a mouse model [51] as well as depressive symptoms in both humans and mice by reducing neuroinflammation [52], ameliorating synaptic dysfunction [53], and promoting neurogenesis [54]. Mice lacking *Pyy* expression also exhibit increased depression-like behavior [55]. Therefore, reduced acetic and butyric acid in BD mice may have influenced mouse behavior through changes in *Pyy* and *Gcg* expression and consequent gut-brain signaling.

Our results also indicate metabolic differences between BD and control mice. BD mice had elevated fecal glucose levels associated with predicted decreased glucose-1-phosphate degradation by gut microbiota. Impaired glucose metabolism has been observed in 50% of individuals with BD [56], and type II diabetes mellitus and BD are comorbid disorders [57]. Elevated sucrose and D-fructose in BD mice further indicate disruptions in gut carbohydrate metabolism that could influence cognitive function [58], signify neurodegeneration [59], and be related to the high comorbidity of metabolic syndromes in BD [60,61]. Metabolic syndromes are also linked to neuroinflammation [62], which could explain increased *Chil3* (chitinase-3-like protein) expression in the amygdalas of BD mice. *Chil3* is produced by myeloid cells in rodents in response to neuroinflammation [63–65], and in humans, elevated serum levels of Chil3L1 (chitinase-3-like protein 1) have been identified in BD patients [66] and individuals with neuroinflammatory conditions [67]. However, there was no differential expression of pro-inflammatory cytokines and glial markers in the colons and amygdalas of BD mice. Furthermore, reduced fecal concentrations of xanthine and hypoxanthine in BD mice, potentially via enhanced inosine 5’-phosphate degradation by gut microbiota, indicate disrupted purine metabolism, which is possibly linked to decreased inosine and upregulated adenosine monophosphate (AMP) and inosinic acid (IMP) in the amygdalas of BD mice. Alterations in purine and adenosine metabolism in the brain are associated with depression [68] and neurodegeneration [59].

An increased abundance of glutamate fermenting *Bacteroides* strains in BD mice could be linked to reduced fecal L-glutamic acid [69], which potentially diminished tetrapyrrole biosynthesis in the gut and lowered levels of the glutamic acid derivative, pyroglutamic acid, in the amygdala. Other studies confirm that gut microbiota can influence pyroglutamic acid levels in the brain [70]. Alterations in amygdalar pyroglutamic acid concentrations in BD mice signify disrupted glutamate metabolism and neurotransmission, which is associated with depression and anxiety in both humans and mice [71,72]. In a piglet model, L-glutamic acid supplementation was also shown to improve small intestinal architecture and influence the expression of amino acid receptors and transporters [73], further connecting reduced fecal L-glutamic acid in BD mice to gut-brain signaling.

Down-regulation glycine, L-tryptophan, uridine, choline, and glycerol in the amygdalas of BD mice further signify altered neurotransmission, neurodegeneration, cellular membrane breakdown, and abnormalities in synaptic plasticity. Glycine reductions are potentially linked to diminished fecal sarcosine, since sarcosine inhibits glycine uptake by glycine-transporter 1 (GlyT1) on neighboring glial cells and can cross the blood-brain barrier [74]. Inhibition of GlyT1 by sarcosine has even been proposed as a treatment for both MDD [75] and schizophrenia [76]. Alterations in sarcosine and glycine production in BD mice could be linked to up-regulation of the S-adenosyl-L-methionine salvage I pathway in the gut. An increase in amygdalar glycine receptor alpha 4 (*Glra4*) expression in BD mice is a potential compensatory mechanism to maximize sensitivity to available glycine in the brain. Given that glycine acts as a co-agonist at N-methyl-D-aspartate (NMDA) receptors, a decline in glycine level could result in abnormalities in synaptic plasticity associated with BD [77]. Glycine supplementation can ameliorate anxiety in rats [78], but elevated glycine in the brain is associated with manic episodes [79], implicating a disrupted glycine homeostasis in BD pathogenesis.

Tryptophan is an established player in gut-brain signaling and both BD and MDD are associated with central and peripheral tryptophan depletion [80–82]. Activation of indoleamine 2,3-dioxygenase (IDO) catabolizes tryptophan; however, because both colonic and amygdalar *Ido-1* expression was similar between BD and control mice, L-tryptophan depletion in the brain was unlikely due to enhanced tryptophan catabolization by IDO. Since Bacteroidota are especially enriched in tryptophan metabolizing strains [83,84], increased Bacteroidota in the guts of BD mice could limit L-tryptophan availability. Fructose malabsorption is also linked to lower serum tryptophan concentrations [85], suggesting elevated gut fructose as a mechanism of tryptophan depletion in BD mice. However, given that fecal L-tryptophan levels were similar between BD and control mice, future work should evaluate disruptions in L-tryptophan brain transportation in BD-FMT mice.

Because uridine, choline, and glycerol are involved in phospholipid biosynthesis, their down-regulation in the amygdalas of BD mice relative to controls indicates disrupted metabolism of cellular membranes, which is linked to the cognitive impairment, reduced neuroplasticity and neurodegeneration associated with BD [86]. Increased Bacteroidota and up-regulated pyrimidine deoxyribonucleosides salvage in the microbiomes of BD mice provide explanation for uridine and glycerol depletion [87,88]. However, reduced brain choline in BD mice is likely independent of choline regulation in the gut since fecal choline levels were similar between BD and control mice. Additionally, there is evidence that reinstitution of uridine and choline could mitigate depression and anxiety symptoms in mice [89,90], and uridine supplementation has been found to significantly increase brain phosphoesters in humans [91].

Our study has several limitations. There is no proof of causality between gut microbiome compositions and gut-brain metabolite profiles and behavior in BD mice. Gut microbiome compositions are inconsistent amongst BD patients, with some studies finding reduced *Faecalibacterium* [92] and Bacteroiodota and increased *Actinobacteria* and *Firmicutes* in BD patients [93]. These inconsistencies are further exacerbated by different regional diets. Additionally, venlafaxine administration was found to elicit microbiome changes in a chronic unpredictable mild stress (CUMS) mouse model; however, *Akkermansia* depletion was still associated with depression-like behaviors [94]. More large-scale microbiome studies of subjects with BD are necessary to fully characterize the BD microbiome and elucidate mechanisms of gut-brain signaling.

Overall, our results demonstrate substantial physiological and behavioral distinctions between mice receiving FMT from a donor with BD and mice receiving FMT from a control donor. BD mice had a greater abundance of Bacteroidota and lower abundances of *P. merdae* and *A. muciniphila*, which are associated with differential fecal SCFA and secondary metabolite production that indicate disrupted carbohydrate metabolism and purinergic signaling. The BD group also displayed increased anxiety-like behaviors and decreased sociability, suggesting that these BD-related behaviors are especially affected by gut-derived signals. Down-regulation of glycine, uridine, choline, glycerol, and L-tryptophan in the amygdalas of BD mice further signifies altered neurotransmission, neurodegeneration, reduced neural plasticity, and cellular membrane breakdown linked to BD. Finally, elevated amygdalar *Chil3* expression in BD mice indicates some neuroinflammatory responses, while no significant differences in the expression of other inflammatory markers were measured between groups. Together, these results suggest a prominent role of gut-brain signaling in our BD-FMT mouse model. Future research should delve deeper into the mechanisms of these gut-brain interactions and perform replication with additional donors. Further exploration of the therapeutic applications of modifying gut microbiota, either through FMT or probiotic interventions, may reveal novel avenues for BD treatment.

## Acknowledgments

This research was funded by “Stadt Graz,” Graz, Austria and a Fulbright-Austrian Marshall Plan Foundation Award to G. B.T. The research of T.M. was supported by Austrian Science Fund (FWF) grants P28854, I3792, DOC-130, and DK-MCD W1226; Austrian Research Promotion Agency (FFG) grants 864690 and 870454; the Integrative Metabolism Research Center Graz; the Austrian Infrastructure Program 2016/2017; the Styrian Government (Zukunftsfonds, doc.fund program); the City of Graz; and BioTechMed-Graz (flagship project).

